# Interactome INSIDER: a multi-scale structural interactome browser for genomic studies

**DOI:** 10.1101/126862

**Authors:** Michael J. Meyer, Juan Felipe Beltrán, Siqi Liang, Robert Fragoza, Aaron Rumack, Jin Liang, Xiaomu Wei, Haiyuan Yu

## Abstract

Protein interactions underlie nearly all known cellular function, making knowledge of their binding conformations paramount to understanding the physical workings of the cell. Studying binding conformations has allowed scientists to explore some of the mechanistic underpinnings of disease caused by disruption of protein interactions. However, since experimentally determined interaction structures are only available for a small fraction of the known interactome such inquiry has largely excluded functional genomic studies of the human interactome and broad observations of the inner workings of disease. Here we present Interactome INSIDER, an information center for genomic studies using the first full-interactome map of human interaction interfaces. We applied a new, unified framework to predict protein interaction interfaces for 184,605 protein interactions with previously unresolved interfaces in human and 7 model organisms, including the entire experimentally determined human binary interactome. We find that predicted interfaces share several known functional properties of interfaces, including an enrichment for disease mutations and recurrent cancer mutations, suggesting their applicability to functional genomic studies. We also performed 2,164 *de novo* mutagenesis experiments and show that mutations of predicted interface residues disrupt interactions at a similar rate to known interface residues and at a much higher rate than mutations outside of predicted interfaces. To spur functional genomic studies in the human interactome, Interactome INSIDER (http://interactomeinsider.yulab.org) allows users to explore known population variants, disease mutations, and somatic cancer mutations, or upload their own set of mutations to find enrichment at the level of protein domains, residues, and 3D atomic clustering in known and predicted interaction interfaces.

## INTRODUCTION

Protein-protein interactions facilitate much of known cellular function. Recent efforts to experimentally determine protein interactomes in human^1^ and model organisms^2-4^, in addition to literature curation of small-scale interaction assays^5^, have dramatically increased the scale of known interactome networks. Studies of these interactomes have allowed researchers to elucidate how modes of evolution affect the functional fates of paralogs^4^ and to examine on a genomic scale network interconnectivities that determine cellular functions and disease states^6^.

While simply knowing which proteins interact with each other provides valuable information to spur functional studies, far more specific hypotheses can be tested if the spatial contacts of interacting proteins are known^7^. In the study of human disease, it has been demonstrated that mutations tend to localize to interaction interfaces and mutations on the same protein may cause clinically distinct diseases by disrupting interactions with different partners^6,8^. However, the binding topologies of interacting proteins (i.e. the relative positions of all atoms in an interaction interface) can only be absolutely determined through resource-intensive X-ray crystallography, NMR, and more recently cryo-EM^9^ experiments, severely limiting the number of interactions with resolved interaction interfaces.

In order to study protein function on a genomic scale, especially as it relates to human disease, a similarly large-scale set of protein interaction interfaces is needed. Thus far, computational methods, such as docking^10^ and homology modeling^11^, have been employed to predict the atomic-level bound conformations of interactions whose experimental structures have not yet been determined. Though it is capable of producing high quality interaction models^12^, docking remains highly specialized and docked models are not yet available on a large scale. Homology modeling has been used to produce models on a large scale^13^, but is only amenable to interactions with structural templates, which comprise <5% of known interactions. Together, co-crystal structures and homology models comprise the currently available precalculated sources of structural interactomes, covering only ∼6% of all known interactions (Fig. 1a-b).

**Figure 1.**
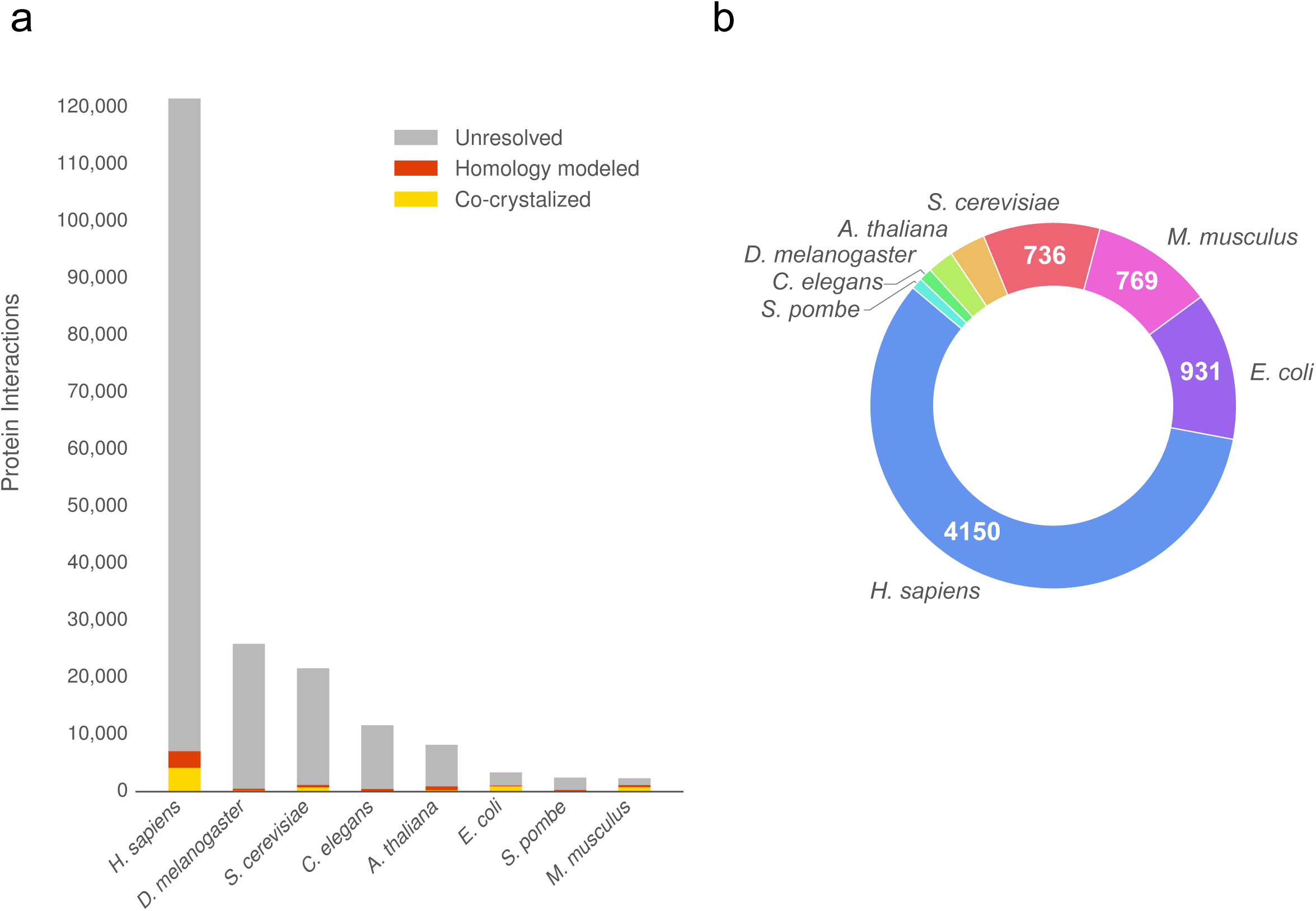
The current size of structural interactomes. (a) The sources of pre-computed structural interactomes and their coverage of known high quality binary interactomes. (b) Interactions from the largest 8 interactomes with experimentally solved structures, which can be used to train a classifier.

While we aim to study disease mutations at atomic-resolution when possible, for the ∼94% of interactions without structural information, a lower-resolution picture of interfaces can provide crucial information for functional studies, and help to complete structural interactome networks to the best of our current capabilities^14^. For instance, residue-level interaction interfaces, where we know which residues are at the interface, but not their precise structural arrangement, can be a great boon to genomic-scale functional analyses^15,16^, and elucidate common modes of human disease^8,17^. Therefore, a multi-scale interactome network containing the highest possible resolution of each protein interaction interface can be an indispensable tool for targeted studies to elucidate pathways and dissect disease mechanisms^4,18^.

Here, we present Interactome INSIDER (<u>**IN**</u>tegrated <u>**S**</u>tructural <u>**I**</u>nteractome and genomic <u>**D**</u>ata brows<u>**ER**</u>), a tool for functional exploration of human disease mutations using the first structurally resolved, multi-scale, proteome-wide human interactome. Interactome INSIDER allows users to find enrichment of disease mutations from popular databases and from user uploads in protein interaction domains, residues, and through atomic 3D clustering in protein interfaces. In order to study disease on a genomic scale, we built an interactome-wide set of protein interaction interfaces by calculating interfaces in experimental co-crystal structures and homology models when available. For the remaining ∼94% of interactions, we applied a new, unified framework, ECLAIR (<u>**E**</u>nsemble <u>**C**</u>lassifier <u>**L**</u>earning <u>**A**</u>lgorithm to predict <u>**I**</u>nterface <u>**R**</u>esidues) to predict the interfaces by applying recent advances in partner-specific interface prediction, such as co-evolution- and docking-based feature construction^19,20^. We used ECLAIR to predict protein interaction interfaces in the full human interactome and for 7 highly studied model organisms (*D. melanogaster, S. cerevisiae, C. elegans, A. thaliana, E. coli, S. pombe*, and *M. musculus*).

Interactome INSIDER (http://interactomeinsider.yulab.org) is deployed as an interactive web server, containing tools for analyzing known and uploaded disease mutations, cancer mutations, and population variants in genome-wide interaction interfaces. Users can also browse predicted interface residues for 184,605 previously un-resolved interactions in human and 7 model organisms, a 15-fold increase over previously known interfaces. Furthermore, for 12,546 interactions with pre-existing sources of structural evidence (co-crystal structures or homology models), we calculate interface residues and display interactive 3D models. Users can search interaction interfaces for enrichment of disease mutations at the level of protein domains, residues, and 3D atomic clustering in a unified interactome composed of all of these sources. We also include relevant functional annotations, such as deleteriousness predictions^21,22^ and biophysical property changes^23,24^ for any proposed mutation or variant that can be viewed in the context of protein and interaction models for a unified functional genomic experience. We anticipate that the marriage of these data sources with our newly predicted full coverage human structural interactome will spur studies of interaction interfaces on a genomic scale.

## RESULTS

In order to build Interactome INSIDER, a tool for genome-wide inference in protein interaction interfaces, we first must construct an interactome-wide set of protein interaction interfaces. Due to lack of structural models, we turned to the well-explored field of protein interaction interface prediction to fill in the gaps in interactomes where neither experimentally-determined co-crystal structures nor homology models are available. While there are well-established methods for predicting protein interactions themselves (i.e. whether or not two proteins interact^)25,26^, we have focused on interactions that have been experimentally determined, but whose interfaces are unknown (**Supplementary Note 1**). For this task, there is a rich literature of methods exploring the potential of many structural, evolutionary, and docking-based methods to predict protein interaction interfaces. However, so far, none of these methods have been used to produce a whole-interactome dataset of protein interaction interfaces (**Supplementary Note 2**).

We created ECLAIR, a unified framework for predicting the interface of any interaction, by leveraging several complementary and proven classification features, including both sequence-based biophysical features, and structural features (**Supplementary Note 3, Supplementary Figs. 1-2**). Furthermore, ECLAIR uses recently proposed features for predicting binding partner specific interfaces, including co-evolutionary^27,28^ and docking-based metrics^20,29^. The advantage ECLAIR offers over previous methods is its ability to be applied to any interaction, while using the most informative set of available interactions for that interaction. In order to accomplish this, ECLAIR is structured as an ensemble of 8 independent classifiers, each covering a common case of feature availability (**Supplementary Notes 4-5, Supplementary Figs. 3-4**). Because each ECLAIR sub-classifier has been trained and tested using a unified set of known protein interaction interfaces, we were able to benchmark each and show that interfaces can be predicted by the single, top-performing sub-classifier that was trained using the full set of features available for each residue (**Supplementary Note 4.2, Supplementary Fig. 5**). In total, we used ECLAIR to predict the interfaces of 184,605 interactions with previously unknown interfaces, including for 114,504 human interactions (**Supplementary Fig. 6**). We supplemented known structural interfaces from co-crystalized proteins and homology models with our predictions to create multi-scale structural interactomes at both the atomic and residue level (Fig. 2a). Finally, in addition to predicting interaction interfaces in 7 model organisms, we created the first multi-scale proteome-wide structural interactome in human for all 121,575 experimentally-determined binary interactions reported in major databases^30-36^ (4,150 with co-crystal structures, 2,921 with homology models, and 114,504 with ECLAIR predicted interfaces; see Materials and Methods), which we used to explore human disease through our new web tool, Interactome INSIDER.

**Figure 2.**
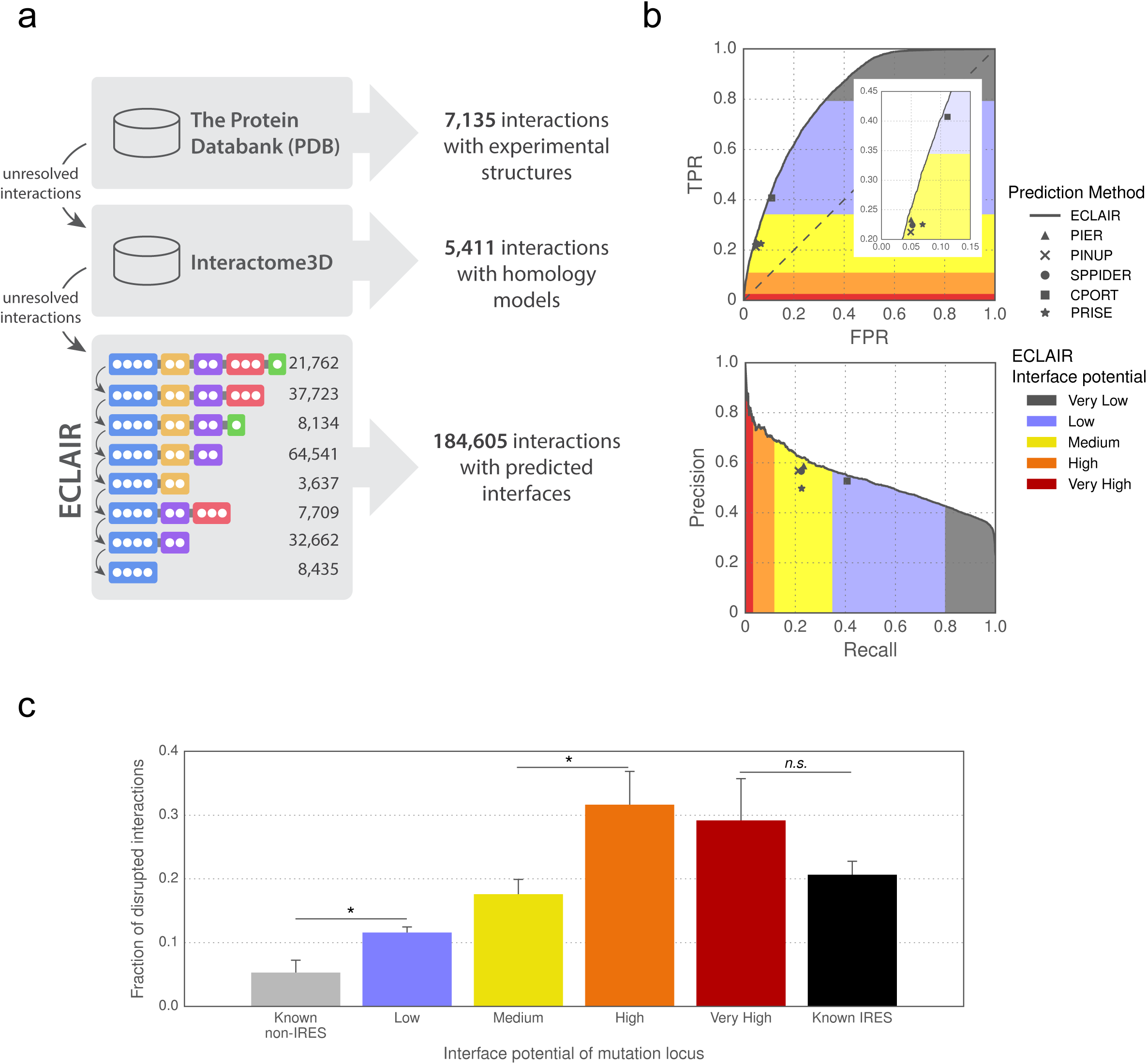
ECLAIR prediction results. (a) Workflow for classifying interfaces for all interactions in 8 species. Interactions without experimentally determined or homology modeled interfaces are classified by ECLAIR. (b) ROC and precision-recall curves comparing ECLAIR with other popular interface residue prediction methods. (c) Fraction of interactions disrupted by the introduction of random population variants in known and predicted interfaces. (* denotes significant (p < 0.05); *n.s*. denotes not significant by a Z-test)

### Comprehensive evaluation of predicted interfaces

In order to use our structural interactomes for functional discovery, we first established that our predictions are of high quality through both machine-learning and biological evaluation. We evaluated the trade-offs between false positive rate and true positive rate, and between precision and recall for each of the 8 independent sub-classifiers that compose ECLAIR (**Supplementary Fig. 5**). As expected, we find that as more informative features are added to subsequent classifiers, the areas under the ROC and precision-recall curves increase, justifying the use of classifiers trained on more features for residues where this information is available.

We next compared ECLAIR to several other prediction methods through two independent validations, in order to establish that ECLAIR’s performance is comparable to other methods. Due to its ensemble nature, we can then apply ECLAIR to many more interactions than would be possible using each of these methods individually. First, we used several readily available predictors^37-41^ to predict interfaces for interactions in our testing set. We find that for the set of interactions for which all classifiers can predict, ECLAIR performs as well or slightly better than these methods by measures of precision, recall, true positive rate and false positive rate (Fig. 2b). Furthermore, for this set of predictors and ECLAIR, we also limited our analyses to only known surface residues, showing that all methods have a slightly lower AUROC (since it is more difficult to distinguish interface from non-interface among only surface residues), however ECLAIR still performs as well or better than all tested methods (**Supplementary Fig. 7**). Finally, we applied ECLAIR to a standard external benchmark set of protein interaction interfaces^42^ which has been used to evaluate the performance of 10 other interface prediction methods^43^. We find that ECLAIR outperforms all benchmarked methods in accuracy, and is comparable to the top performers in all other metrics (**Supplementary Table 1**). Furthermore, ECLAIR is applicable to any interaction, while methods in this benchmark rely on single-protein structure inputs, making them less applicable to genome-wide studies.

We also performed >2,000 mutagenesis experiments to measure the rate at which population variants in our predicted interfaces disrupt interactions compared to variants within known co-crystal interfaces and non-interfaces (see Material and Methods). Since it is known that mutations at protein interfaces are more likely to break interactions^6,18^, we hope to show that mutations in our predicted interfaces also break their corresponding interactions at a significantly higher rate than those known to be away from the interface and at similar rates compared to mutations in known interfaces (it is important to note that only ∼21% of these mutations at known interfaces disrupt corresponding interactions since we tested population variants randomly selected from the Exome Sequencing Project^44^, many of which are believed to be benign). Using our high-throughput mutagenesis yeast two-hybrid assay^18^, we find that the disruption rates for mutations at known interface residues are quite similar to disruption rates for mutations of predicted interface residues (Fig. 2c). Furthermore, even mutations of residues with a ‘Low’ predicted interface potential are significantly more likely to disrupt interactions than mutations of residues known to be away from the interface. This suggests that there is viable functional signal in ECLAIR predictions, as even interfaces predictions in the ‘Low’ potential category show some signs of similar functional properties to known interfaces.

### Interactome INSIDER, a genomics toolbox for interactome studies

We built Interactome INSIDER, a tool for searching for functionally enriched areas of protein interactomes, and for browsing our multi-scale structural interactome networks. Interactome INSIDER contains all 197,151 protein interactions whose interfaces have been either experimentally determined, homology modeled, or predicted using ECLAIR. Specifically for human, Interactome INSIDER contains interface information for all 121,575 experimentally-determined binary interactions reported in major databases^30-36^. Additionally Interactome INSIDER includes 56,159 disease mutations from HGMD^45^ and ClinVar^46^ and analyzed 1,300,352 somatic cancer mutations from COSMIC^47^ to compute their per-disease, pre-calculated enrichment in protein interaction interfaces at the residue level, domain level, and through atomic clustering. Furthermore, the site includes information on >600,000 population variants from the Exome Sequencing Project^44^, 1000 Genomes Project^48^ and more^49^ (see Materials and Methods). Users can then search Interactome INSIDER by protein to retrieve all interaction partners and their interfaces, or by disease to retrieve all interaction interfaces that are enriched for mutations of that disease. Additionally, users can upload their own set of mutations to find how they are distributed in the interactome and whether they are enriched in any protein interaction interfaces at the residue, domain, and atomic levels (Fig. 3).

**Figure 3.**
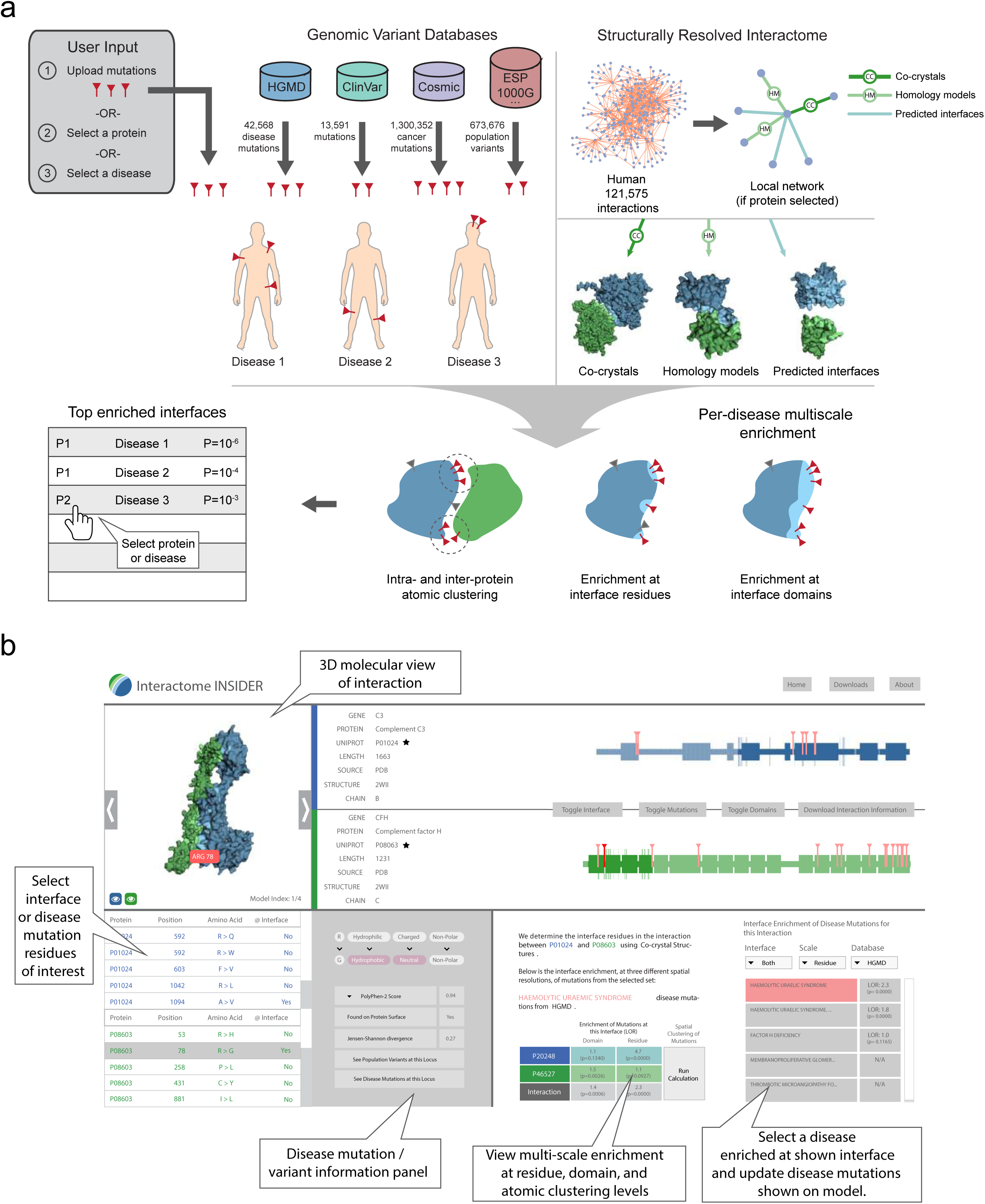
Flowchart showing the sources and computational workflow for calculating mutation and variant enrichment using the Interactome INSIDER web interface. Users may submit their own mutations or select sets of known disease and cancer mutations to assess their enrichment in interface domains and residues, or compute 3D atomic clusters of mutations in proteins and across interfaces.

Since our goal is to use Interactome INSIDER to explore protein function, especially disease, in interactomes, we next investigated the functional biological properties of our predicted interaction interfaces. These studies involve measuring functional properties of our de novo predicted interfaces (those without prior experimental evidence) and comparing these measurements to those of known interfaces from co-crystal structures. Importantly, these known properties of interaction interfaces are completely separate of the features used for training ECLAIR, and thus provide an unbiased and functionally relevant means to assess the utility of our predicted interfaces. Showing that our predicted interfaces have many of the same functional properties as known interfaces suggests their applicability to functional genomic studies.

Many studies have probed the link between interactome networks and disease^50,51^, and it is well established that disease mutations are enriched at structural interfaces of interacting proteins^6,8,52^, suggesting that disruption of binding with one or more partners may contribute to the disease phenotype. Though not all disease mutations will appear at the interfaces of interactions, and can act via other mechanisms, such as destabilizing proteins entirely^18^, their enrichment at interfaces is a statistically significant global trend^6,8^. However, >40% of known missense and nonsense human disease mutations cause alterations to proteins lacking any structurally resolved interaction interfaces. To test whether our predicted interfaces may be useful for the study of disease, and thus help address this knowledge gap, we looked at their positions relative to disease mutations. We find that disease mutations also preferentially occur in our predicted interfaces, at similar rates to known interface residues occurring in PDB co-crystal structures (Fig. 4a), indicating the viability of using predicted interfaces to study molecular disease mechanisms. Furthermore, each more confident bin of predicted interface residues is more likely to contain disease mutations than the previous, showing that ECLAIR prediction scores are correlated with true protein function.

**Figure 4.**
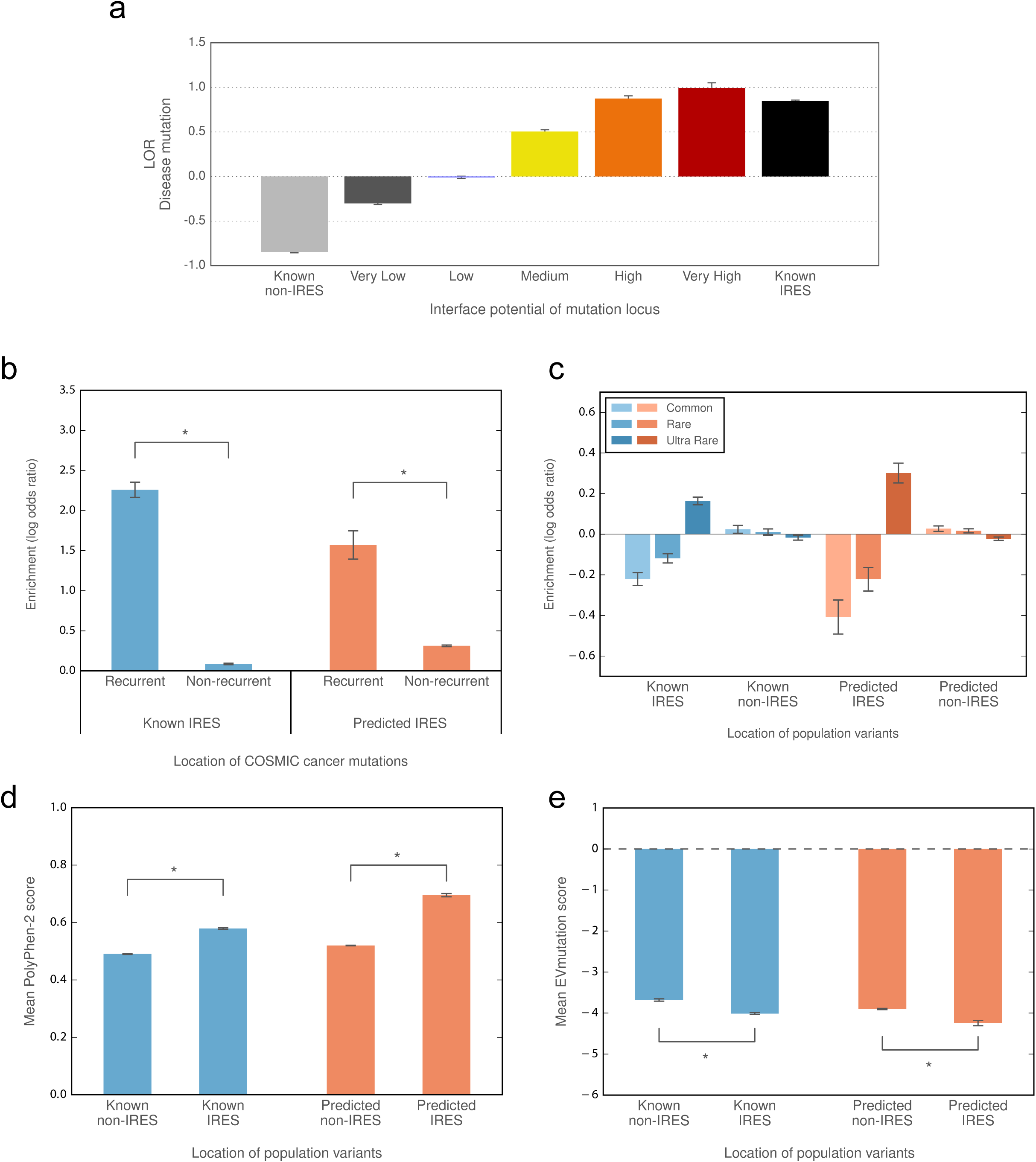
Functional properties of predicted interfaces. (a) Enrichment of disease mutations in predicted and known interfaces. (b) Enrichment of recurrent cancer mutations in predicted and known interfaces. (c) Enrichment of rare and common population variants in predicted and known interfaces. (d) Predicted deleteriousness of population variants in known and predicted interfaces (using PolyPhen-2). (e) Predicted effects of population variants in known and predicted interfaces (using EVmutation). (* denotes significant, p < 0.05 by a Z-test)

Similarly, we looked at the locations of somatic cancer mutations from COSMIC in our interface-resolved human interactome. We specifically focused on recurrent cancer mutations as these are known to be more likely than infrequently observed mutations to be functional drivers^53-55^. We find a marked enrichment of recurrent cancer mutations in our predicted interfaces compared to outside our predicted interfaces (Fig. 4b). Furthermore, the same trend is observed inside and outside of known interfaces from co-crystal structures, suggesting that the functional links between cancer and the potential disruption of protein interactions can be observed within our entire human interface dataset. We also looked at the distribution of population variants, and show that their placement in and out of predicted interfaces matches that of known interfaces, with rarer mutations showing an enrichment in protein interfaces (Fig. 4c). Furthermore, we show that population variants in our predicted interfaces are more likely to be damaging to protein function than variants outside of predicted interfaces, as predicted by PolyPhen-2^21^ (Fig. 4d) and EVmutation^56^ (Fig. 4e), matching the established trend for experimentally determined interfaces^57^.

These enrichment analyses and the matched results between our predicted interfaces and known interfaces in co-crystal structures further confirm the validity of ECLAIR predictions. More importantly, users can take advantage of these enrichment analyses through our Interactome INSIDER web server to better dissect large-scale whole-genome and whole-exome sequencing datasets to help identify novel disease-associated genes and mutations. For instance, if known disease mutations are significantly enriched in a specific interaction interface, this information could be used to complement and further boost the confidence of patient and disease-specific variants in the same interface that have been prioritized by other methods (e.g., co-segregation^58^, mutation burden^59^). This can be particularly helpful to sieve through the large number of variants of unknown significance generated by large-scale sequencing studies. Furthermore, if known disease mutations are enriched in a specific interface of a protein whose involvement in the disease is already understood, this could still suggest its interaction partner’s mechanistic involvement in the disease through this specific interface, even if the partner is not yet known to be associated with the disease.

To illustrate the usefulness of Interactome INSIDER, we searched for sub-networks in the human interactome that are enriched for disease mutations associated with a single disease by calculating the enrichment of disease mutations in interaction interfaces interactome-wide, a functionality also available to users via the Interactome INSIDER website. This allowed us to identify the TGF-β/BMP signaling pathway, which is known to be involved in juvenile polyposis syndrome (JPS)^60^, and contains multiple proteins harboring JPS mutations (Fig. 5a). We focused on a specific group of mutations in the SMAD4-SMAD8 interface, which can be found using 3D atomic clustering. Using our mutagenesis Y2H assay, we were able to test a JPS mutation (SMAD4 Y353S)^61^, which is at the interface of SMAD4-SMAD8, and show that it breaks this interaction, implicating SMAD8 in JPS. Although SMAD8 (also known as SMAD9) has not been reported to harbor JPS mutations in HGMD^45^, its involvement in the disease has been suggested^62^, showing the ability of Interactome INSIDER to implicate new proteins in disease. Furthermore, Y353S is not predicted by ECLAIR to be at the interface of SMAD4 and another of its binding partners, RASSF4, and indeed, through our Y2H experiment, does not break this interaction, demonstrating the functional insight Interactome INSIDER can provide about differential interfaces and how they might be relevant to understanding the molecular mechanisms of disease.

**Figure 5.**
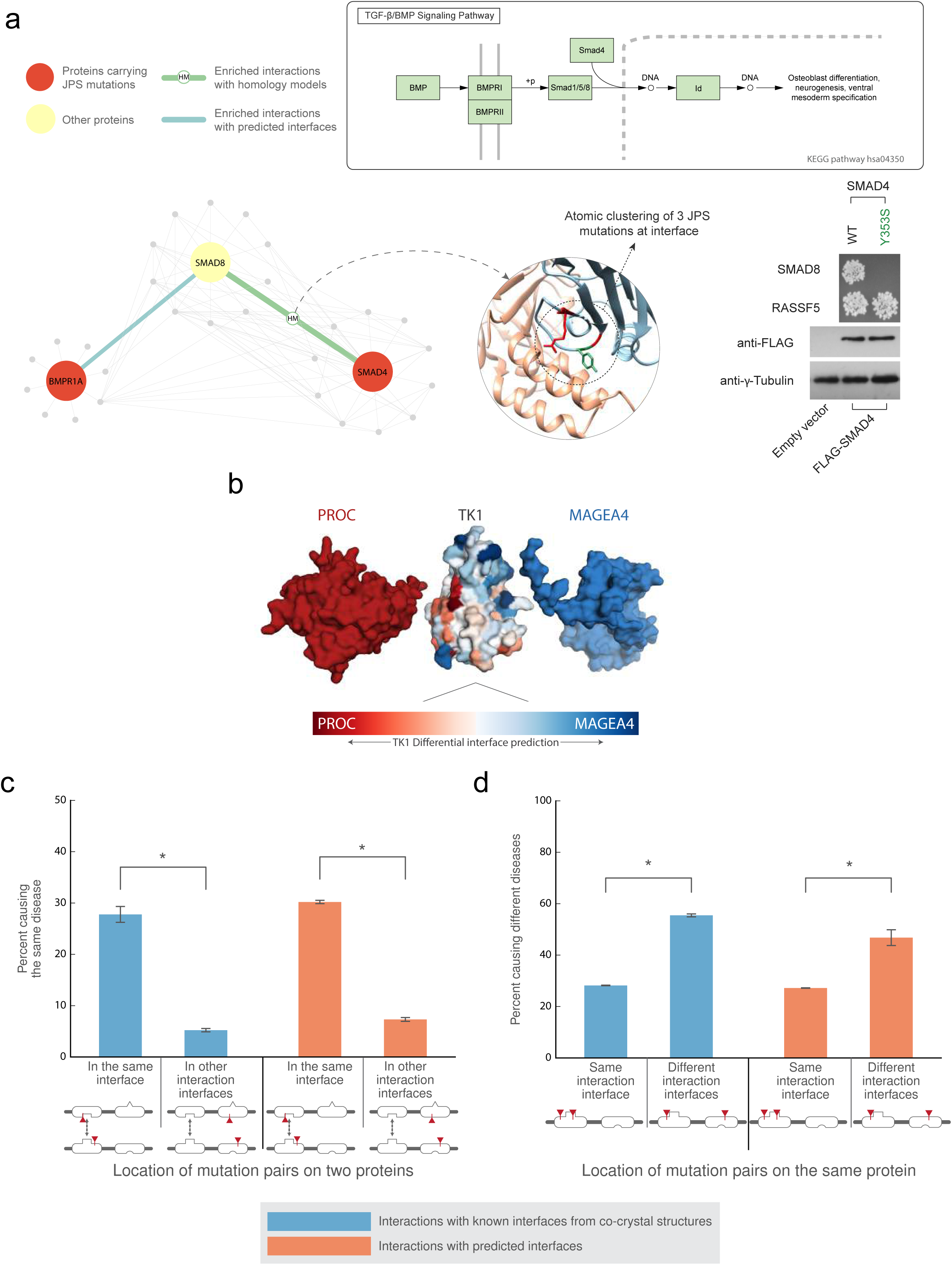
Interaction partner specific interface prediction. (a) The TGF-β/BMP signaling pathway. Atomic clustering reveals a mutation hotspot for juvenile polyposis syndrome at the interface of SMAD8 and SMAD4. One of these mutations (Y353S) on SMAD4 is confirmed to selectively break the interaction with SMAD8. This mutation is not predicted by ECLAIR to be at the interface of SMAD4-RASSF5, and is shown to leave this interaction intact. (b) Superimposed docking results of two different partners with TK1. The differentially predicted interfaces of TK1 with each of its partners corresponds with the orientation of the docked poses. (c) Pairs of disease mutations across interaction interfaces tend to cause the same disease more often than pairs of mutations in other interfaces of the same proteins with different partners. (d) Pairs of disease mutations in different interfaces of the same protein tend to cause different diseases more often than pairs of mutations in the same interface. (* denotes significant, p < 0.05 by a Z-test)

### Disease etiology revealed by partner-specific interfaces

In addition to providing full coverage of interfaces in the human interactome, one major benefit that Interactome INSIDER provides is the ability to interrogate different interfaces for the same protein dependent upon its binding partner. For the study of protein function and disease, this is especially important as a protein may maintain different functional pathways through different interfaces, and disruption of one interface may leave another intact^4,8^. To demonstrate the potential of Interactome INSIDER to tease apart interface-specific disease mutation etiologies, where the same mutation can cause differential effects with two different binding partners, we first investigated an example of differential interface prediction using ECLAIR. Here we highlight an interaction whose predicted interfaces are strongly influenced by a single partner-specific feature, molecular docking. In Figure 5b, the protein TK1 is shown colored by its docked pose with each of two partners, PROC and MAGEA4. We note that the predicted interface residues on TK1 are drastically different for each partner, and that the areas with elevated interface potential correspond to the position of the two docking results. Even though these interaction interfaces were predicted using features additional to docking, this demonstrates how even a single partner-specific feature can lead to differential interface predictions.

The ability of Interactome INSIDER to reveal interaction partner-specific effects can also be demonstrated as a global trend in our ECLAIR-predicted interfaces. As discussed, this is important because disruptions of different interfaces of the same protein may cause differential disease states; for instance, disruption of one interface may cause a disease while disruption of another may not cause any detriment to protein function. To demonstrate this on a large-scale, we looked at pairs of disease mutations in the human interactome that appear at interaction interfaces. It has been shown that pairs of disease mutations in interacting proteins cause the same disease when located in the interaction interface more often than mutations located in interaction interfaces with separate partners^8^. We performed the same analysis using differential interaction interfaces predicted by ECLAIR and find the same trend—mutation pairs in the interface of two interacting proteins are much more likely to cause the same disease than mutation pairs in other interfaces of the same proteins that do not mediate the given interaction (Fig. 5c). We also find that mutation pairs on the same protein, but in separate interfaces with different binding partners tend to cause different diseases (Fig. 5d). Moreover, this trend is observed in both known and predicted interfaces. This shows that ECLAIR is able to use partner-specific features such as docking and co-evolution to predict different interfaces depending on the binding partner of a protein that match established trends of pleiotropy and locus heterogeneity in known interfaces^8^. Importantly, this indicates that Interactome INSIDER can be used to form functional hypotheses about the specificity of mutations to specific interactions and molecular pathways.

Using Interactome INSIDER to find sub-networks in the human interactome enriched for disease mutations associated with a single disease, we also uncover a set of interacting proteins known to harbor mutations causal of hypertrophic cardiomyopathy (HCM)^63^, a disease marked by enlargement of the myocardium heart muscle that can become fatal, and automatically recapitulate the core constituents of a known KEGG pathway related to the same disease (Fig. 6). These proteins were identified by enrichment of disease mutations in their shared interaction interfaces and, in the case of TNNI3-TNNC1, using cross-interface atomic clustering of disease mutation positions in 3D. Access to enrichment and 3D atomic clustering tools for this disease and users’ uploaded mutations is available via the Interactome INSDIER web interface.

**Figure 6.**
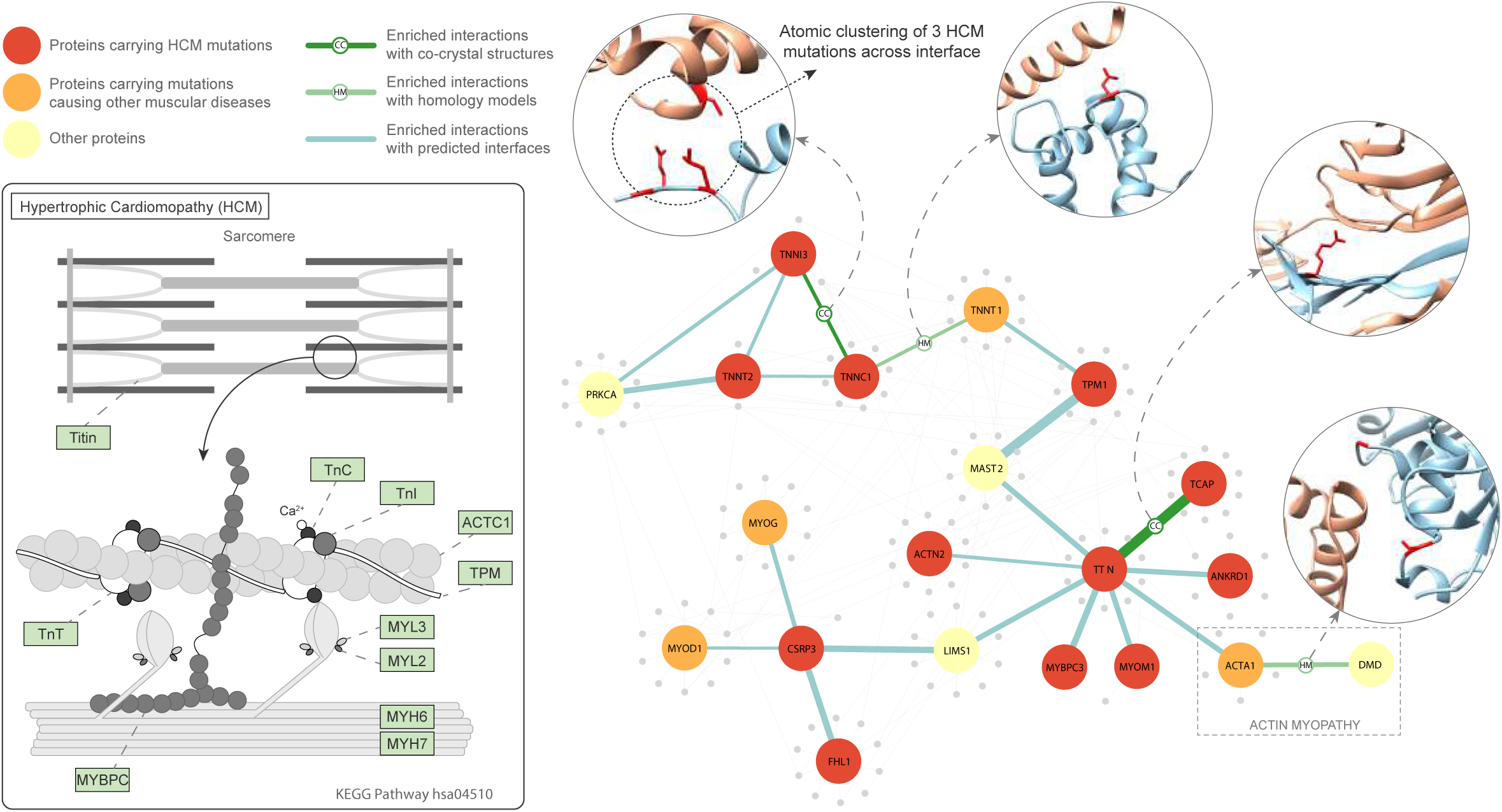
The hypertrophic cardiomyopathy pathway. Interactome INSIDER readily discovers the core components of this KEGG pathway known to be involved in the disease hypertrophic cardiomyopathy (HCM). Interfaces are noted for their enrichment of HCM mutations or, in the case of TNNI3-TNNC1, the presence of 3D atomic clusters. Through enrichment analyses, Interactome INSIDER also reveals an additional protein, CSRP3, which is not part of the known KEGG pathway, but is enriched for HCM mutations at predicted interfaces with known pathway members. Furthermore, TTN’s interface with ACTA1 is enriched for HCM mutations, but a separate interface of ACTA1 with its binding partner dystrophin is enriched with mutations causing a distinct disorder, actin myopathy, demonstrating Interactome INSIDER’s ability to discover cases of differentiable function mirroring differential interfaces.

Interestingly, in addition to identifying known members of the HCM pathway, Interactome INSIDER also identified several additional proteins, including CSRP3, MYOM1, ANKRD and TCAP, which are not part of the known KEGG pathway, but carry HCM mutations enriched at their respective interaction interfaces with members of the pathway. We also identify a protein, TNNT1, which, although it contains no HCM mutations of its own, can be implicated in HCM by interacting with two proteins TPM1 and TNNC1, which are enriched for HCM mutations at their interfaces with TNNT1. Finally, we note that Interactome INSIDER reveals cases of partner-specific interfaces in this pathway. For instance, the known HCM pathway protein TTN’s interface with ACTA1 is enriched for HCM mutations, and ACTA1 mutations are increasingly linked to HCM^64,65^. On the other hand, a separate interface of ACTA1 with its binding partner dystrophin is enriched with mutations causing a distinct disorder, actin myopathy^66^. This shows how ACTA1 can play roles in two different diseases through separate interaction interfaces with TTN and dystrophin, and demonstrates Interactome INSIDER’s unique ability to discover such cases of differentiable function mirroring differential interfaces.

## DISCUSSION

Interactome INSIDER is an integrative information center for genomics studies in the structural human interactome. By leveraging several sources of protein interaction interfaces, including experimentally determined co-crystal structures, homology models, and predicted interfaces, Interactome INSIDER allows scientists to probe for functional insights in whole interactomes, and to predict disease etiologies based on network topology and specific structural interfaces at several scales of resolution. Our new interface prediction pipeline, ECLAIR, incorporates many previously validated strategies and features for predicting protein interaction interfaces in whole genomes, allowing Interactome INSIDER to be the first resource to show that predicted interfaces can be used for functional analyses in whole interactomes, especially for the study of human disease.

We anticipate Interactome INSIDER will help to bridge the divide between genomic-scale datasets and structural proteomic analyses, both now and in the future. Now that large-scale sequencing data from many contexts are readily available, for instance from whole-genome/whole-exome population variant studies^44,67^ and cancer studies^68,69^, researchers have become increasingly interested in ways to assess the potential functional consequences of variants on a genomic scale^59,70-72^. For instance, recently we and others have developed methods to predict functional cancer driver mutations by finding hotspots of mutations in the structural proteome^52,54,73^. With the comprehensive map of protein interfaces presented, we can now go a step further to predict specific etiologies of cancer and disease based on induced biophysical effects^74,75^ that may break interactions. Because our interface map is partner-specific, it can also be applied to predict pleiotropic effects, wherein several mutations in a single protein may affect different pathways depending upon which binding interfaces are mutated^8^. This could be the basis for designing new therapeutics and for rational drug design to selectively target specific protein functional sites^76^.

The scale of interactomes and functional genomic data in Interactome INSIDER uniquely enables it to be useful for genomic studies. While at least one previous resource, dSysMap^77^, is able to display disease mutations in structural interfaces, it is limited to 9,875 human interactions with either co-crystal structures or homology models, severely limiting its applicability to genomic studies by offering the same small slice of the interactome that has been studied extensively. Interactome INSIDER on the other hand contains interfaces for an additional 111,700 human interactions alone, which have never been available before in any repository. Furthermore, unlike dSysMap, Interactome INSIDER contains somatic cancer mutations, population variants, and mutation functional annotations, as well as interfaces for 7 model organisms with potential use in model systems studies, which have proven useful in the study of human disease^78^ and for studying molecular evolution^4,79^. Thus, we intend Interactome INSIDER be a more broadly applicable resource, with the ability to inform many aspects of genomic studies, from identifying functional regions of proteins, to incorporating orthogonal information about known mutations and functional effects in these regions.

With future increases to the scale of biological databases from which we derive features, we expect that Interactome INSIDER will come to encompass even higher confidence predictions for many more interactions, thereby becoming increasingly applicable to functional studies. This may also address some limitations of structural databases today. For instance, the PDB is depleted of disordered proteins^80^, and it has been shown that disordered regions can form interfaces^81^. Since ECLAIR has not been trained on disordered interfaces, it is unlikely to predict new disordered interfaces. However, the ensemble classifier structure of ECLAIR uniquely positions it to incorporate all newly-available evidence into interface predictions without sacrificing quality or scale, ensuring the highest quality map of interaction interfaces now and in the future. Furthermore, the addition of new variants, especially cancer mutations and population variants from large-scale sequencing studies, will only increase the value of performing systems-level explorations with Interactome INSIDER.

## MATERIALS & METHODS

### Interaction datasets

We compiled binary protein interactions available for *H. sapiens, D. melanogaster, S. cerevisiae, C. elegans, A. thaliana, E. coli, S. pombe*, and *M. musculus* from 7 primary interaction databases. These databases include IMEx^82^ partners DIP^30^, IntAct^31^, and MINT^32^, IMEx observer BioGRID^33^, and additional sources iRefWeb^34^, HPRD^35^, and MIPS^36^. Furthermore, iRefWeb combines interaction data from BIND^83^, CORUM^84^, MPact^85^, OPHID^86^, and MPPI^87^. We filtered these interactions using the PSI-MI^88^ evidence codes of assays that can determine experimental binary interactions (**Supplementary Table 2**), as these are interactions where proteins are known to share a direct binding interface that we can then predict^5^. In total, we curated 197,151 interactions in these 8 species including the full experimentally determined binary interactome in human (121,575 interactions) (**Supplementary Note 1**). Those interactions with known interface residues based on available co-crystal structures in the Protein Data Bank (PDB)^89^ were set aside for use in training and testing the classifier. Interactions without known interface residues comprise the set for which we make predictions.

### Testing and training sets for interface residue prediction

For those interactions with known co-crystal structures in the PDB, we calculate interface residues for their specific binding partners. To identify UniProt protein sequences in the PDB, we use SIFTS^90^, which provides a mapping of PDB-indexed residues to UniProt-indexed residues^49^. For each interaction and representative co-crystal structure, interface residues are calculated by assessing the change in solvent accessible surface area of the proteins in complex and apart using NACCESS^91^. Any residue that is at the surface of a protein (≥15% exposed surface) and whose solvent accessible surface area (SASA) decreases by ≥1.0 Å^2^ in complex is considered to be at the interface. We aggregate interface residues across all available structures in the PDB for a given interaction, wherein a residue is considered to be at the interface of the interaction if it has been calculated to be at the interface in one or more co-crystal structures of that interaction (all other residues are considered to be away from the interface). In building our final training and testing sets, we only consider interactions for which aggregated co-crystal structures have combined to cover at least 50% of UniProt residues for both interacting proteins.

The training and testing sets each include a random selection of 400 interactions with known co-crystal structures, of which 200 are heterodimers and 200 are homodimers (**Supplementary Table 3**). To ensure an unbiased performance evaluation, we disallowed any homologous interactions (i.e. interactions whose structures could be used as templates for homology modeling) between the training and testing set. We also disallowed repeated proteins between the two sets to avoid simply reporting a remembered shared interface between a protein and multiple binding partners, thereby artificially elevating the performance of our classifier on the testing set.

### Hyperparameter optimization with TPE

In order to train our ensemble of classifiers that comprise ECLAIR, we used the tree-structured Parzen estimator approach (TPE)^92^, a Bayesian method for optimizing hyperparameters for machine learning algorithms. TPE models the probability distribution *p*(*x|y*) of hyperparameters given evaluated loss from a defined objective function, *L(x)*. We selected the following loss function to minimize based on classical hyperparameter inputs and residue window sizes:

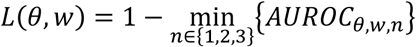
where *x* is comprised of *θ*, a set of hyperparameters, and *w*, a set of residue window sizes. The evaluation metric, *AUROC_n_*, is the area under the roc curve for the n^th^ left-out evaluation fold in a three-fold cross-validation scheme. We then used TPE to randomly sample an initial uniform distribution of each of our hyperparameters and window sizes and evaluate the loss function for each random set of inputs. TPE then replaces this initial distribution with a new distribution built on the results from regions of the sampled distribution that minimize *L(x)*:

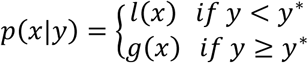
where *y^*^* is a quantile γ of the observed y values so that *p(y* < *y^*^)* = γ. Importantly, *y^*^* is guaranteed to be greater than the minimum observed loss, so that some points are used to build *l(x)*. TPE then chooses candidate hyperparameters to sample as those representing the greatest expected improvement, *EI*, according to the expression:

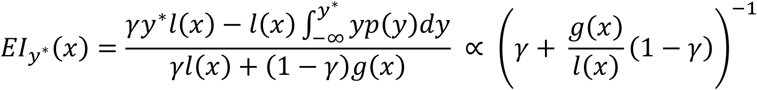
In order to maximize *EI*, the algorithm picks points *x* with high probability under *l(x)* and low probability under *g(x)*. Each iteration of the algorithm returns *x*^*^, the next set of hyperparameters to sample, with the greatest *EI* based on previously sampled points.

### Training the classifier

The ECLAIR classifier was trained in three stages, using a custom wrapper of the scikit-learn^93^ random forest^94^ classifier to allow for use of TPE to search both algorithm hyperparameters and residue window sizes simultaneously. In all cross-validations performed, we allowed TPE to search the following hyperparameters, beginning with uniform distributions of the indicated ranges: (1) minimum samples per leaf (0-1000), (2) maximum fraction of features per tree (0-1), and (3) split criterion (entropy or gini diversity index). The number of estimators (decision trees) in each random forest was fixed at either 200 for training the feature selection classifiers, or 500 for training the full ensemble. We also allowed TPE to search over residue window sizes (± 0-5 residues for a total window of up to 11 residues, centered on the residue of interest). This was achieved by allowing extra features for neighboring residues to be included at the time of algorithm initialization.

In the first stage of training, cross-validation using TPE was performed on classifiers trained using only features from 1 of the 5 feature categories. The feature or set of features from each category with the minimum loss was selected to represent that category in building the ensemble classifier (**Supplementary Table 4**). In the second stage, the ensemble classifier was built of 8 random forest classifiers, each trained on different subsets of feature categories, and hyperparameters and window sizes were again chosen using cross-validation and TPE (**Supplementary Table 5**). In the final stage, following performance measurement on the testing set, the 8 sub-classifiers were retrained using the full set of 3,447 interactions with at least 50% UniProt residue coverage in the PDB, using the same hyperparameters and window sizes found in the previous step.

### Evaluating the ensemble

After training and optimizing using only the training set, we predicted interface residues in a completely orthogonal testing set. For each sub-classifier of the ensemble, all residues in the testing set that could be predicted (given the full set of necessary features or a superset) were ranked according to their raw prediction scores to produce ROC and precision-recall plots.

### Benchmarking against other methods

Interfaces for interactions in our testing set were computed using several popular interface prediction methods^37-41^. We compiled a set of representative protein structures from the PDB for each protein in our testing set, selecting the structure with the highest UniProt residue content based on SIFTS and excluding any PDB structures of interacting protein pairs from our testing set. We then evaluated the precision, recall, and false positive rate for proteins that were able to be classified by all methods. These represent point estimates of these metrics for the external methods with binary prediction scores.

We also compared ECLAIR to 10 popular methods for interface prediction by predicting interfaces in a standard benchmark set of protein complexes^42^ (**Supplementary Table 1**). Here, we followed the experimental procedures laid out by Maheshwari *et al.^43^*, and excluded complexes in which the receptor is <50 or >600 amino acids, where the interface is made up of <20 residues, or where multiple interfaces are present.

### Predicting new interfaces

We retrained the ensemble using all available co-crystal structures, including those from both testing and training sets, a standard machine learning practice that makes maximal use of labeled data^95^. Using this fully trained ensemble of classifiers, we predicted interface residues for the remaining 184,605 interactions not resolved by either PDB structures or homology models. Sub-classifiers were ordered based on the number and information content of features used in their training. Each residue was then predicted by only the top ranking classifier of the ensemble trained on the full set or a subset of available features for that residue.

### Interface enrichment and 3D atomic clustering

Interface domain enrichment, residue enrichment, and 3D atomic clustering can be calculated through the Interactome INSIDER web interface. For enrichments presented in this study, we accessed all disease mutations from the Human Gene Mutation Database (HGMD)^45^ and ClinVar^46^, recurrent cancer mutations appearing ≥ 6 times in COSMIC^47^, and population variants from the Exome Sequencing Project^44^ to compute the log odds ratio:

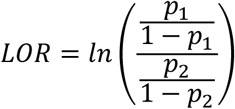
where *p_1_* is the probability of a mutation or variant being at the interface and *p_2_* is the probability of any residue being at the interface. We computed the log odds ratio for residues in each of the interface prediction potential categories. We also computed the log odds ratio for interactions with known interfaces from PDB co-crystal structures, defined as all known interface residues from NACCESS calculations and all residues in Pfam^96^ domains with ≥ 5 interface residues. For the disease mutation enrichment analysis (Fig. 4a, we used all disease mutations available from HGMD, and the following numbers of mutations occurred in each category: 10,196 Very Low, 10,547 Low, 2,970 Medium, 1,135 High, and 305 Very High. We also computed enrichment of 18,638 mutations in known interfaces and 17,760 mutations in known non-interfaces (from co-crystal structure evidence).

To perform 3D atomic clustering of amino acid loci of interest, we used an established method^54^ for clustering and empirical *p*-value calculation and applied it to multi-protein clustering, wherein clusters can occur across an interaction interface. Here, we perform complete-linkage clustering^97^ in the shared 3D space of both proteins, and iteratively, and randomly rearrange mutations in each protein to produce an empirical null distribution of cluster sizes.

### Mutagenesis validation experiments

We performed mutagenesis experiments in which we introduced random human population variants from the Exome Sequencing Project^44^ into known and predicted interfaces. We randomly selected mutations of predicted interface residues in each of the top four ECLAIR categories (Low – Very High). As positive and negative controls, we also selected random mutations of known interface and non-interface residues in co-crystal structures in the PDB. The selected mutations were then introduced into the proteins according to our previously published Clone-seq pipeline^18^ and their impact (either disrupting or maintaining the interaction) was assessed using our yeast two-hybrid assay (**Supplementary Note 6**). In this manner, we tested the impact of 2,164 mutations: 1,664 in our predicted interfaces and 500 in known interface and non-interface residues from co-crystal structures. In Figure 2c, we report the fraction of tested interface residue mutations that caused a disruption of the given interaction for each of the interface residue bins.

### Web server

Interactome INSIDER is deployed as an interactive web server (http://interactomeinsider.yulab.org) containing known and predicted interfaces for 197,151 protein interactions in 8 species, as well as variants and functional annotations mapped relative to the residues in the human proteome. For each interaction, the most reliable, high-resolution model is presented, i.e. co-crystal structures are always displayed in lieu of homology models, and all remaining unresolved interactions are predicted by our ECLAIR classifier. Co-crystal structures are derived from the PDB, with extraneous chains removed for each interaction, and homology models are computed by MODELLER^11^ and downloaded from Interactome3D^13^. For both types of structural model, we computed all residues at the interface over all available models, and allow users to view any model from which a unique interface residue has been calculated. For predicted interfaces, a non-redundant set of single protein models are shown when available, with locations of predicted interface residues indicated. In total, the resource contains 7,135 interactions with co-crystal structures, 5,411 with homology models, and 184,605 with predicted interfaces.

Interactome INSIDER also includes pre-calculated enrichment of mutations derived from several sources: 56,159 disease mutations from HGMD^45^ and ClinVar^46^ and 1,300,352 somatic cancer mutations from COSMIC^47^. It also includes 194,396 population variants from the 1000 Genomes Project^48^, 425,115 from the Exome Sequencing Project^44^, and 54,165 catalogued by UniProt^49^. Predictions of deleteriousness for all variants and any user-submitted variants within the curated interactomes are obtained from PolyPhen-2^21^ and SIFT^22^, and biophysical property change guides (i.e. polar to non-polar, hydrophobic to hydrophilic) are also displayed for convenience. Mutation and variant enrichment analyses can be triggered by the user for existing variants or for user-submitted sets within interacting protein domains, residues, and 3D clustering using the atomic coordinates of structures when available.

